# QTL analysis of floral morphology polymorphisms between *Gilia yorkii* and *G. capitata*

**DOI:** 10.1101/2023.11.22.568340

**Authors:** Joseph M DeTemple, Daniel H Chitwood, Veronica Mosquera, Clinton J Whipple

## Abstract

Speciation is a complex process typically accompanied by significant genetic and morphological differences between sister populations. In plants, this can result in divergent floral morphologies causing pollinator differences and reproductive isolation between populations. Here, we explore floral trait differences between two recently diverged species, *Gilia yorkii* and *G. capitata*. The distributions of floral traits in parental, F1, and F2 populations are compared, and groups of correlated traits are identified. We describe the genetic architecture of floral traits through a quantitative trait locus (QTL) analysis using an F2 population of 187 individuals. While all identified QTLs were of moderate (10-25%) effect, interestingly, many QTL intervals colocalized on Chromosomes 6 and 9, suggesting that sets of traits may share a common genetic basis. Our results provide a framework for future identification of genes involved in the evolution of floral morphology.

## Introduction

The color and dimensions of floral organs vary naturally within plant populations, facilitating adaptive change to different selective pressures (Sapir *et al*. 2021). Divergence in floral traits (e.g. color and size of petals, anther and style lengths, and throat length) makes important contributions to reproductive isolation and speciation (Rieseberg *et al*. 2006). Divergent floral morphologies can arise from pollinator-driven selection (Wessinger *et al*. 2014; Brothers *et al*. 2013; Chen *et al*. 2020; Campitelli *et al*. 2018; Kostyun *et al*. 2019), breeding system incompatibilities (Kostyun *et al*. 2019; Roux and Pannell 2019; Mertens *et al*. 2018; Eckert *et al*. 1996; Goodwillie *et al*. 2006), and random genetic drift (Roux and Pannell 2019; Tremblay and Ackerman 2001; Yoshida *et al*. 2008). Despite the central role floral morphology plays in plant evolution and taxonomy, surprisingly little is known about the genetic mechanisms that underlie the morphological evolution of flowers.

In order to understand the genetic architecture of heritable, divergent floral morphologies, Quantitative Trait Locus (QTL) analysis of floral traits can be applied to hybrid populations. This approach uses recombination events resulting from a cross of parents that differ in heritable traits of interest to correlate phenotypes with genotypes and determine the number and effect size of loci regulating those traits (Miles and Wayne 2008). High levels of genetic and morphological variation are ideal for QTL mapping, and can be obtained by crossing highly divergent parental lines. Within plants, wide crosses between morphologically distinct species often produce hybrids, such as in grasses (Freeling 2001), *Mimulus* (Bradshaw *et al*. 1995), *Aquilegia* (Hodges *et al*. 2002), and *Epidendrum* (Pinheiro *et al*. 2010). A common negative trade-off to these wide crosses is reduced fertility of the hybrid, which compromises the ability to create mapping populations. However, when fertile hybrid progeny can be produced, fine-mapping of QTL has provided valuable insights into the genes and polymorphisms that underlie morphological evolution (Ballerini *et al*. 2020; Liang *et al*. 2023).

In this study, we report a fertile inter-specific cross within the *Gilia* genus that facilitates the mapping of divergent floral morphologies. The leafy-stemmed gilias (*Gilia* section *Gilia*, Polemoniaceae) comprise 11 species found in North and South America. They are annual plants with small white or purple flowers and a raceme or panicle inflorescence (Grant 1966) (Porter 2012). Hybridization between (and within) leafy-stemmed gilia species was explored by Verne Grant in the 1950s (Grant 1949), revealing weak to strong barriers to reproduction existing between species and, in some cases, between populations of the same species (Grant 1966). Subsequent to the initial biosystematic work of Grant, a new leafy-stemmed gilia species, *G. yorkii*, was discovered in 1998 (Shevock and Day 1998). A molecular phylogeny showed that *G. yorkii* is closely related to *G. capitata* (Johnson and Porter 2017). Previously, we showed that certain accessions of *G. capitata* produce fertile hybrids when crossed to *G. yorkii* (Jarvis *et al*. 2022), opening the door for an interspecies QTL analysis of floral traits. *G. yorkii* and *G. capitata* differ in numerous floral traits including flower color (Jarvis *et al*. 2022), size, stamen exsertion, and pedicel length. Both species are diploid annuals and have simple growth requirements, making them convenient for genetic analysis. Because of their ease of cultivation, crossing compatibility, and divergent morphology, *G. yorkii* and *G. capitata* represent an ideal system to probe genetic causes of inter-species variation in floral traits.

Here we report floral morphology QTLs that distinguish *G. yorkii* and *G. capitata* by creating an interspecific F2 mapping population. We find 23 QTLs linked to 17 different floral traits, many of which have overlapping intervals, suggesting a common genetic basis for these traits. This study adds to the growing body of literature documenting QTLs for floral trait differences between species and provides a framework for future identification of the underlying genes.

## Materials and methods

With the exception of plants grown for initial measurements of floral traits in the parent lines (see below) all Plant Materials, Mapping Populations, DNA isolation, Sequencing, Genome Assembly, Annotation, Genotyping-by-Sequencing (GBS), Genetic Map, and QTL analysis were all performed as outlined in (Jarvis *et al*. 2022).

### Plant Growth Conditions

*G. yorkii* and *G. capitata* plants were grown indoors in a growth room prior to growing the mapping population to get initial trait measurements. These plants were grown in Sungro soilless potting mix supplemented with 18g/l osmocote in 6-in pots under 16-hour days using fluorescent lights, with a constant temperature of 20°C.

### Floral Traits

Floral traits were measured digitally from images of dissected, fresh flowers. Flowers were cut horizontally at the base to separate the calyx, corolla, and ovary from the pedicel. The calyx was slit from a sinus to the base. The corolla was slit from the base of the corolla tube to the sinus of petal lobes, taking care to be on one side of the free filament of the corresponding stamen. Corolla and calyx were laid on a glass microscope slide coated with double-sided tape. Images were taken with a Leica s8apo dissecting scope. All images were then processed using the Leica Application Suite X (LASX) software using the measuring tool, and all measurements were recorded in an Excel spreadsheet. Measured floral traits were divided into four categories: corolla, reproductive, sepal, and other traits. Corolla traits include Petal length (PeL), Petal lobe length (PeLL), Petal lobe width (PeLW), Petal tube length (PeTL), Petal tube width (PeTW), Throat length (TrL), Filament length (FL), and Free filament length (FFL). Reproductive traits include Anther length (AL), Anther width (AW), Style length (SyL), Stigma length (SgL), and Ovary shape (OS). Sepal traits include Sepal length (SeL), Sepal sinus length (SeSL), Sepal tooth length (SeTL), and Sepal midrib width (SeMW). Other traits include Pedicel length (PdL), Internode length (IL), Days to flower (DTF), and Vegetative rosette diameter (VRD). A full list of all traits is found in Table 1, and traits will be referred to by their abbreviations. A diagram including the major floral parts is found in Figure 1.

**Table 1.**
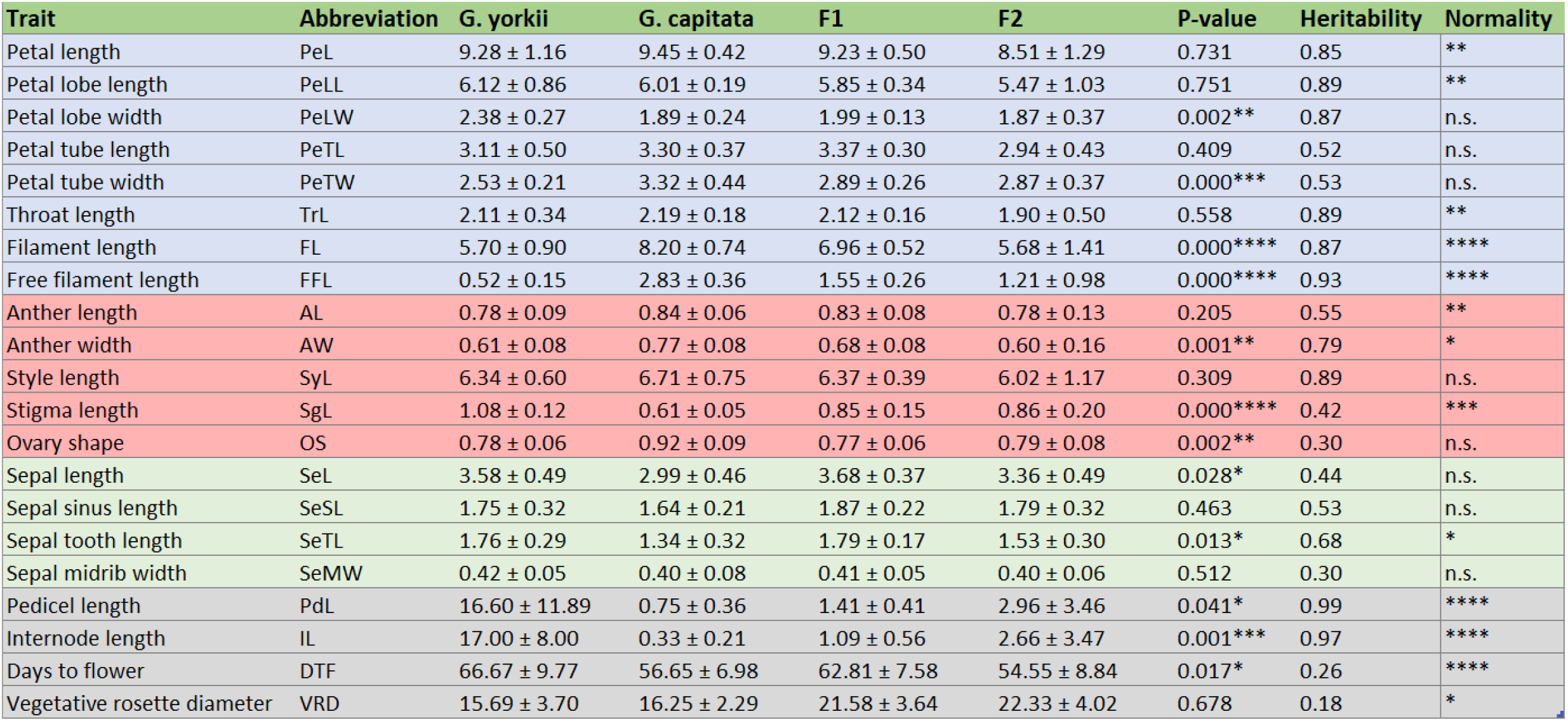
Mean trait values and standard errors for *G. yorkii, G. capitata*, F1, and F2 populations, T-test p-values between parent values, Broad-sense Heritability (*H*^2^), and Shapiro-Wilks Normality test of the distribution (* = 0.1 significance level; ** = 0.01; *** = 0.001; **** = 0.0001; n.s. means the distribution is not significantly different from normal). Blue, red, green, and gray row colors correspond to corolla, reproductive, sepal, and other traits respectively. All measurements are in millimeters (mm).

**Figure 1.**
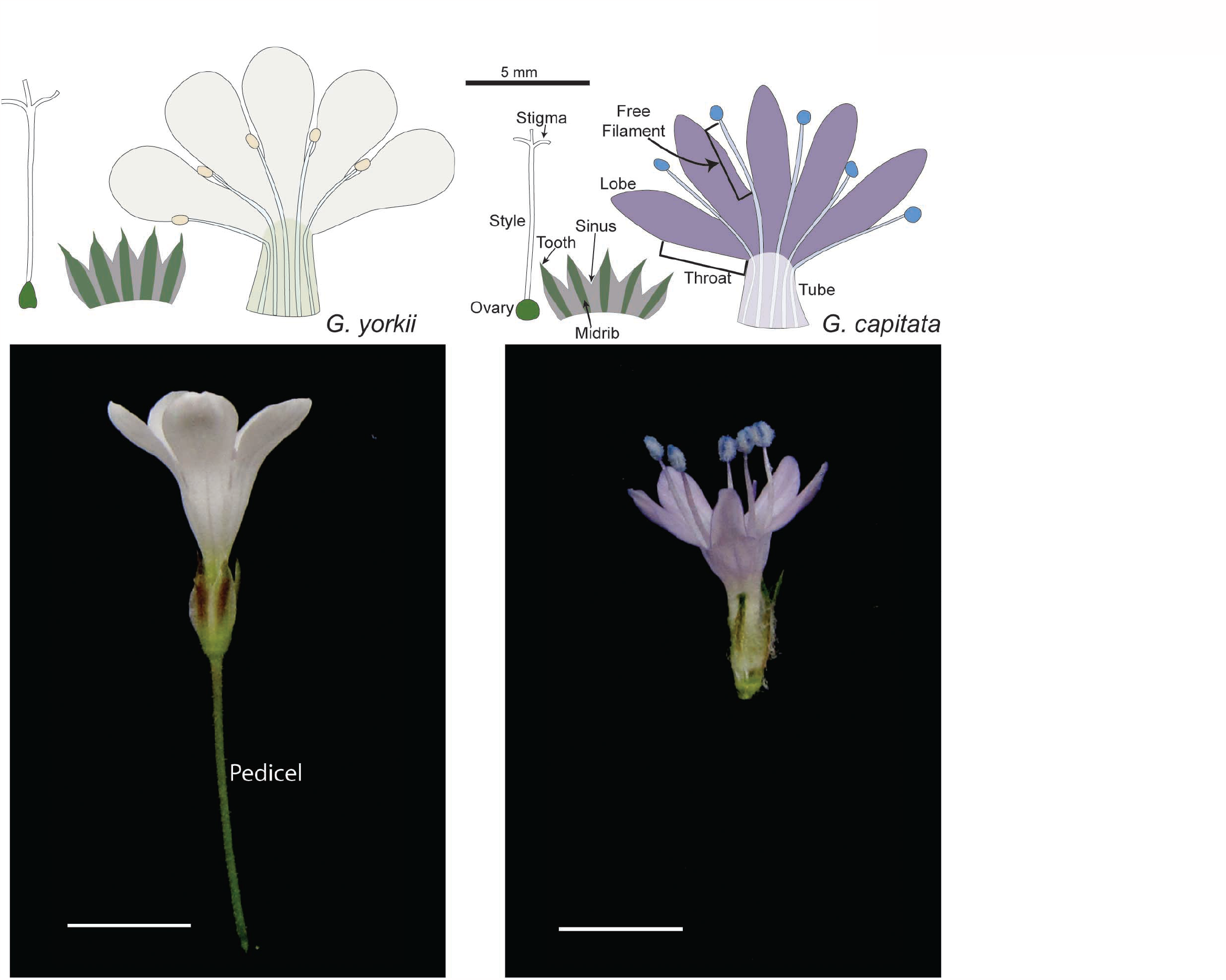
Typical flowers of *G. yorkii* (left) and *G. capitata* (right). Floral parts corresponding to traits included in the QTL analysis are labeled.

**Figure 2.**
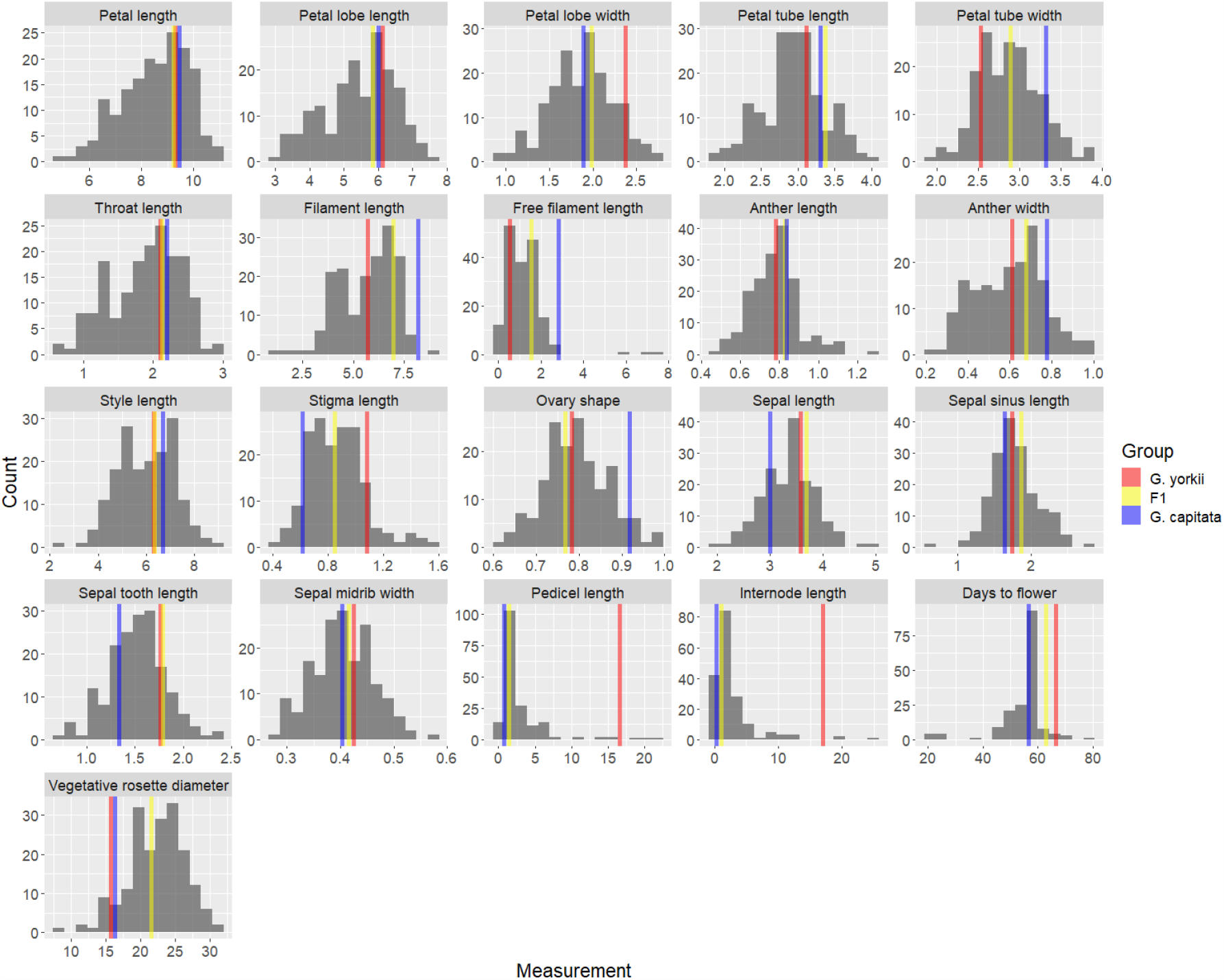
Floral trait distributions in the F2 generation. G. yorkii, F1, and G. capitata population means are shown in red, yellow, and blue lines respectively. Axis scales are millimeters (mm).

Data for the Procrustes analysis (see below) of floral shape were collected by placing landmark points at the corners of the opened floral tube, the sinus of each petal lobe, two points at the widest part of each petal lobe, and one point at the tip of the petal lobe.

### Statistical Analysis

Mean values, standard error, and Student’s t-test between parent values were all calculated in R using base functions. Broad-sense heritability (*H*^2^), or the proportion of variance not due to environmental factors alone, was calculated manually in Excel using the formula:

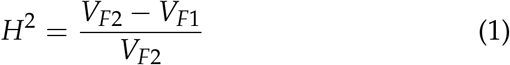

where *V*_*F*1_ and *V*_*F*2_ are the variances of the F1 and F2 populations respectively.

Trait histograms were generated in R using the “ggplot2” package. Correlations between traits were calculated using Pearson’s correlations in R, and the correlation plot was generated using the “corrplot” package. Procrustes Analysis and subsequent Principal Components were generated using the “procGPA” function from the “shapes” package. QTL data was loaded and analyzed in the “rqtl” package (Broman *et al*. 2003; Broman and Sen 2009). LOD significance level for QTL markers were determined by a permutation test of 100,000 replications across all traits yielded a 95% level of 4.16 LOD score. QTL intervals were declared using the “lodint” function using a LOD drop of 1 and the position of the qtl index corresponding to the marker with the highest LOD in intervals that extended above the significance threshold.

## Results

### Floral morphology polymorphisms between *G. capitata* and *G. yorkii*

The overall floral morphology of *G. capitata* and *G. yorkii* is similar and shared with diverse *Gilia* species and many related Polemon-icaceae (Figure 1). The sepals are united at the base with a free toothed apex and a distinct green (occasionally infused with purple) band running along the midrib flanked by a light-green to hyaline margin. Like the sepals, petals of both species are also fused basally, and this fused region can be divided into a narrow basal tube which transitions into a flaring throat. Distal to the throat, five unfused lobes are symmetrically arranged. Five stamens alternate with the petals, and the stamen filaments are adnate with the basal petal along both the tube and throat, and then extend slightly on a free filament between petal lobes. The pistil is comprised of a distinct round to oval-shaped ovary at the base and an elongated style which branches apically into three distinct stigmas.

Within this shared floral groundplan, multiple differences across *G. capitata* and *G. yorkii* are apparent. *G. capitata* flowers are notably smaller with exserted stamens, narrower petal lobes and a hardly perceptible pedicel compared to the larger *G. yorkii* flowers with inserted stamens, wide petal lobes, and a long pedicel. To identify consistent morphological differences between the parent species, we dissected flowers and measured multiple traits associated with sepal, petal, stamen and carpel morphology on 30 flowers sampled from (5-6) individuals of each species grown in uniform growth room conditions (Table 2). Significant differences were found in overall length of sepals (SeL), the free sepal tooth length (STL), and the sepal midrib width (SMW) with *G. yorkii* being the larger species. Similarly, in the petal whorl *G. yorkii* had significantly larger petal length (PeL), petal lobe length (PeLL), tube length and throat length, as well as petal lobe width (PeLW). The petal tube width (PeTW) was also significantly different, but in this case *G. capitata* was larger. In the stamen whorl, total filament, free filament length (FFL) and anther Width (AW) were all significantly larger in *G. capitata*. In the pistil whorl, the ratio of ovary width:length, a measure of ovary shape (OS), was significantly larger in *G. capitata*. While there was no significant difference in style length, stigmas were significantly longer in *G. yorkii*. Most traits, with the exception of SgL, OS, SeL, SeMW, DTF, and VRD, had relatively high heritabilities (*H*2 > 0.5), showing that most floral traits are good candidates for QTL analysis.

**Table 2.** Mean trait values and standard errors for *G. yorkii* and *G. capitata* populations grown in a growth chamber, and T-test p-values between parent values. Blue, red, and green row colors correspond to corolla, reproductive, and sepal traits respectively. All measurements are in millimeters (mm).

| Trait | Abbreviation | G. yorkii | G. capitata | P-value |
| --- | --- | --- | --- | --- |
| Petal length | PeL | 10.53 ± 0.38 | 9.61 ± 0.79 | 0.000 |
| Petal lobe length | PeLL | 3.35 ± 0.27 | 2.81 ± 0.51 | 0.001 |
| Petal lobe width | PeLW | 2.41 ± 0.19 | 3.46 ± 0.20 | 0.000 |
| Petal tube length | PeTL | 2.57 ± 0.21 | 2.43 ± 0.51 | 0.000 |
| Petal tube width | PeTW | 6.84 ± 0.23 | 6.44 ± 0.33 | 0.000 |
| Throat length | TrL | 2.99 ± 0.18 | 1.80 ± 0.32 | 0.040 |
| Filament length | FL | 6.68 ± 0.21 | 7.53 ± 1.30 | 0.001 |
| Free filament length | FFL | 0.93 ± 0.16 | 2.44 ± 0.62 | 0.000 |
| Anther length | AL | 0.62 ± 0.04 | 0.64 ± 0.05 | 0.211 |
| Anther width | AW | 0.46 ± 0.03 | 0.56 ± 0.05 | 0.000 |
| Style length | SyL | 2.92 ± 0.30 | 2.68 ± 0.97 | 0.185 |
| Stigma length | SgL | 1.50 ± 0.11 | 1.23 ± 0.17 | 0.000 |
| Ovary shape | OS | 1.34 ± 0.05 | 1.37 ± 0.13 | 0.000 |
| Sepal length | SeL | 0.40 ± 0.34 | 0.35 ± 0.39 | 0.014 |
| Sepal sinus length | SeSL | 7.27 ± 0.23 | 7.02 ± 0.23 | 0.000 |
| Sepal tooth length | SeTL | 1.19 ± 0.18 | 0.63 ± 0.21 | 0.599 |
| Sepal midrib width | SeMW | 0.67 ± 0.04 | 0.94 ± 0.06 | 0.000 |

For most traits measured, the variance was larger in *G. capitata* than in *G. yorkii*. While *G. yorkii* is self-compatible and a highly inbred line was used for all measurements, the self-incompatible DeTemple et al. 3 *G. capitata* was sib-crossed for 5 generations, and significantly less inbred than *G. yorkii*. It is possible that some of the additional variance in the *G. capitata* floral traits are a result of residual segregating genetic variation.

### Distribution of Floral traits in an F2 mapping population

We generated an F2 mapping population of 189 individuals as described previously (Jarvis *et al*. 2022), and grew these in a greenhouse along with 18 *G. capitata* individuals, 12 *G. yorkii* individuals, and 12 F1 hybrid individuals. All floral traits described above were measured across these plants (Table 1). In addition, we measured pedicel length (PdL), inflorescence internode length (IL, the distance between the subterminal flower and the next lowest node), and vegetative traits including days to flowering (DTF), and vegetative rosette diameter (VRD). Unexpectedly, petal length measurements (PeL, PeLL, PeTL, and TrL) that were significantly different in growth room plants were no longer significant in the greenhouse environment, which may suggest a G × E interaction for these traits.

Normality of the floral traits in the F2 population was measured using the Shapiro-Wilk test. Of the 19 floral traits, 10 show normality at the *α* = 0.01 level. This includes PeLW, PeTL, PeTW, AW, SeL, SeSL, SeTL, SeMW, OS, and SyL. For the other 9 traits, upon visual inspection it was determined that FFL, IL, and PdL had the most severely skewed distributions, and log transformation was performed before QTL analysis (Goh and Yap 2009). Of these skewed distributions, FFL was the only trait where *G. capitata* had a higher mean value than *G. yorkii*.

Broad-sense heritability of the floral traits ranged between 0.30 to 0.99. While many of the floral traits had high heritability values (greater than 0.6), some traits had lower heritabilities. In particular, PeTL, PeTW, SeSL, and SeMW have heritabilites ranging from 0.30 to 0.53 (Table 1). It is not immediately apparent why these traits relating to the petal tube and calyx have low heritabilities compared to the other traits, although it does limit their potential as traits to follow up on in further genetic analyses. Two other traits, DTF and VRD, show extremely low heritability values of 0.26 and 0.18 respectively. These traits are expected to be highly sensitive to environmental conditions, and this is confirmed by the observation that the F1 and F2 populations have similar variances of these traits. The most puzzling traits with relatively low heritability were AL and SgL: these traits are both related to reproduction and occupy very prominent positions within the flower, but do not seem to be under the same genetic control as other adjacent floral elements, with heritability values of 0.55 and 0.42 respectively.

### Trait Correlations

To better understand how individual floral traits are connected, we calculated Spearman’s Correlations for all floral traits and identified groups of correlated traits. Correlations between all floral traits are shown in Figure 3. The largest group of correlated traits consisted of length and width dimensions of the petals, stamens and style (PeL, PeLL, PeLW, FL, AW, and SyL), all of which showed strong positive correlations with each other. Unexpectedly, anther 4 Interspecies floral morphology QTL in *Gilia* length did not strongly correlate with this entire group but did correlate with anther width. In addition, several petal and stamen width and length traits (PeTL, PeTW, and FFL) correlated with one or more, but not all, traits in this group. Calyx measurements (SeL, SeSL, SeTL, and SMW) comprise a second distinct set of correlations from the petal-stamen-style group. Correlations of all the traits in this group with sepal length were strongest, while correlations between sepal midrib width, sepal tooth length, and sepal sinus length by themselves were low to moderate. Internode elongation in the inflorescence was strongly correlated with pedicel length (Figure 3). VRD and DTF did not correlate significantly with any floral morphological traits.

**Figure 3.**
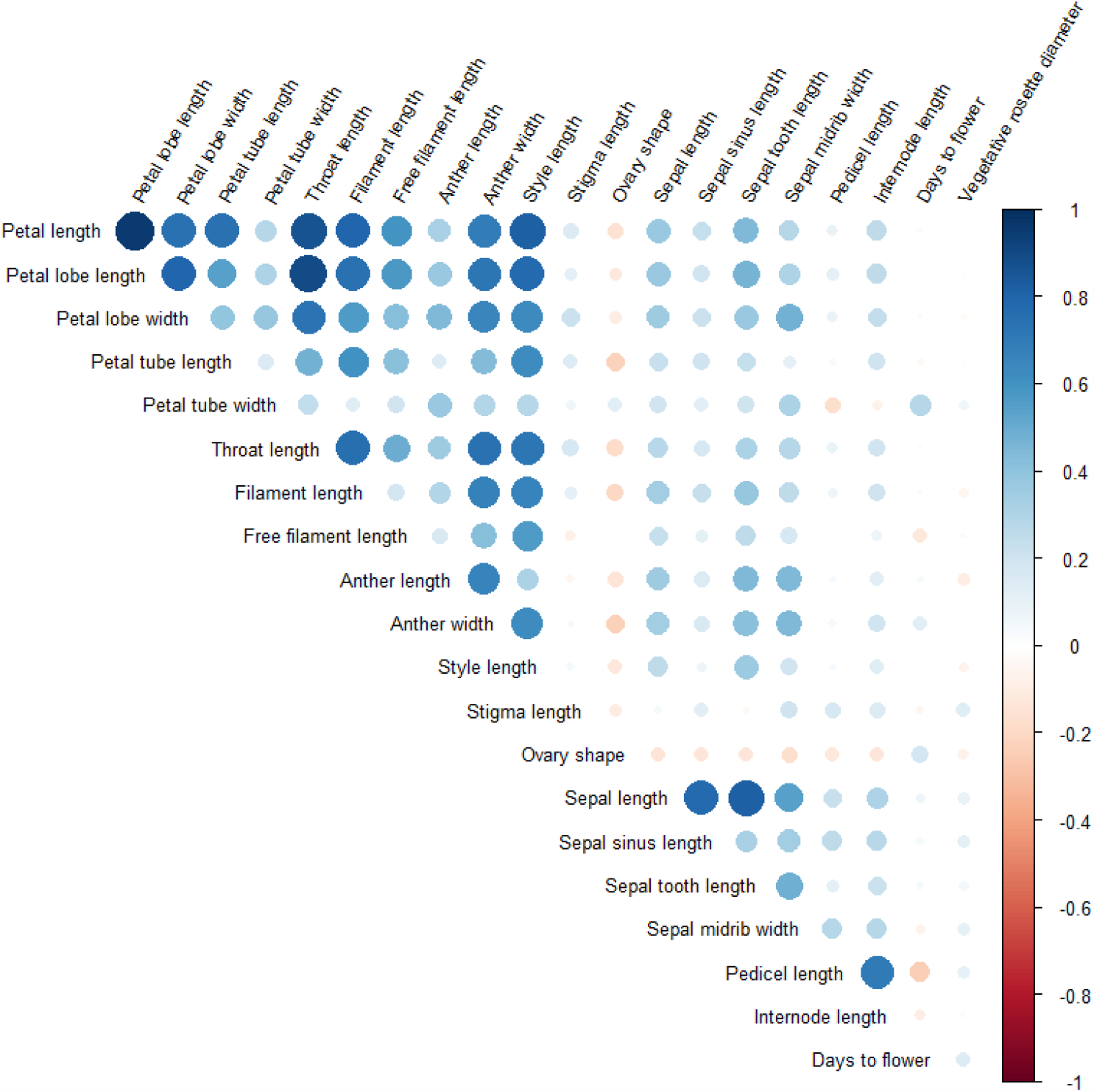
Correlations between floral traits. Color represents the direction of correlation, whereas intensity and circle size represent the degree of correlation.

### Morphometric Analysis

To investigate whether changes in overall floral morphology are significantly different between the parent populations, we collected landmark data points from whole flowers for a Procrustes analysis, which uses a Principal Component Analysis (PCA) to reduce the dimensionality of the data. The first principal component (Figure 4) accounts for 30.9% of the variance present in the combined DeTemple et al. 5 parent population. Visually, PC1 captures much of the variation we observed previously between *G. yorkii* and *G. capitata* flowers, where *G. yorkii* has wider petal lobes and a longer floral tube as 6 Interspecies floral morphology QTL in *Gilia* compared to *G. capitata*, and it effectively separates *G. yorkii* and *G. capitata* individuals. When F2 landmark data are transformed into this principal component background, they generate intermediate values (Figure 5). QTL mapping of the F2 PC1 values resulted in no significant QTLs, suggesting that the genetic basis of overall shape is highly quantitative in nature and major effect loci for this trait are not present in our QTL analysis.

**Figure 4.**
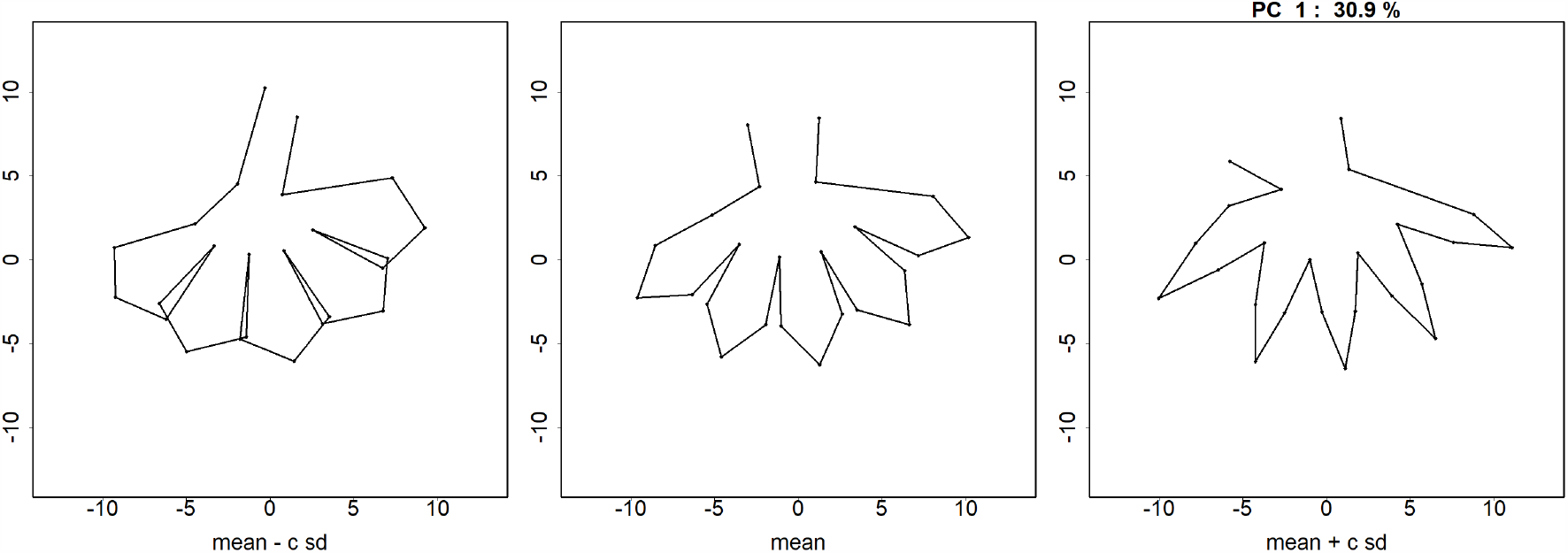
Principal Component 1 from PCA analysis of Procrustes-adjusted landmark data.

**Figure 5.**
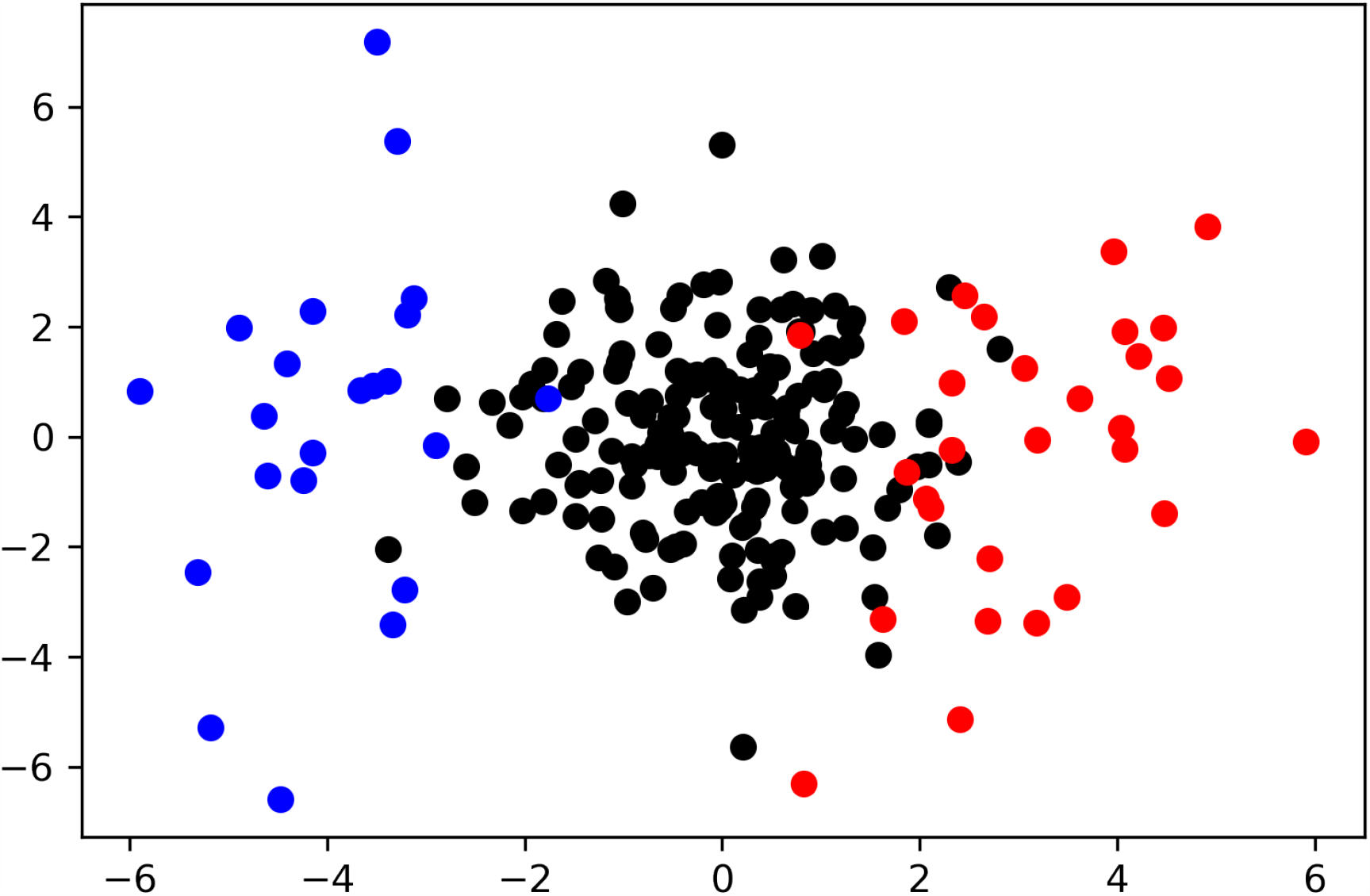
F2 population projected onto PC1 (x-axis), PC2 (y-axis) space of parent populations. Each dot represents an individual plant, and the red, blue, and black colors represent *G. yorkii, G. capitata*, and F2 individuals respectively.

### QTL Analysis

To perform trait mapping, a genetic map was created from SNPs identified from the Genotyping-by-Sequence data. After filtering for several marker quality attributes (see methods, (Jarvis *et al*. 2022)), a final map consisting of 5,335 markers was used for QTL mapping of floral traits in the F2 hybrid population of 189 individuals. Mapping was performed using an extended Haley-Knott regression in the ‘rqtl’ package. Intervals were calculated by a LOD drop of 1.5 from the highest single marker. A total of 23 significant QTL were identified across all floral traits. QTLs were found on six out of the nine chromosomes present in *Gilia* (Fig. 6, and Table 3). Thirteen traits had one significant QTL each, three traits (PeLL, PeLW, and SyL) mapped to two QTL each, and four QTL on different chromosomes were identified for SgL.

**Table 3.**
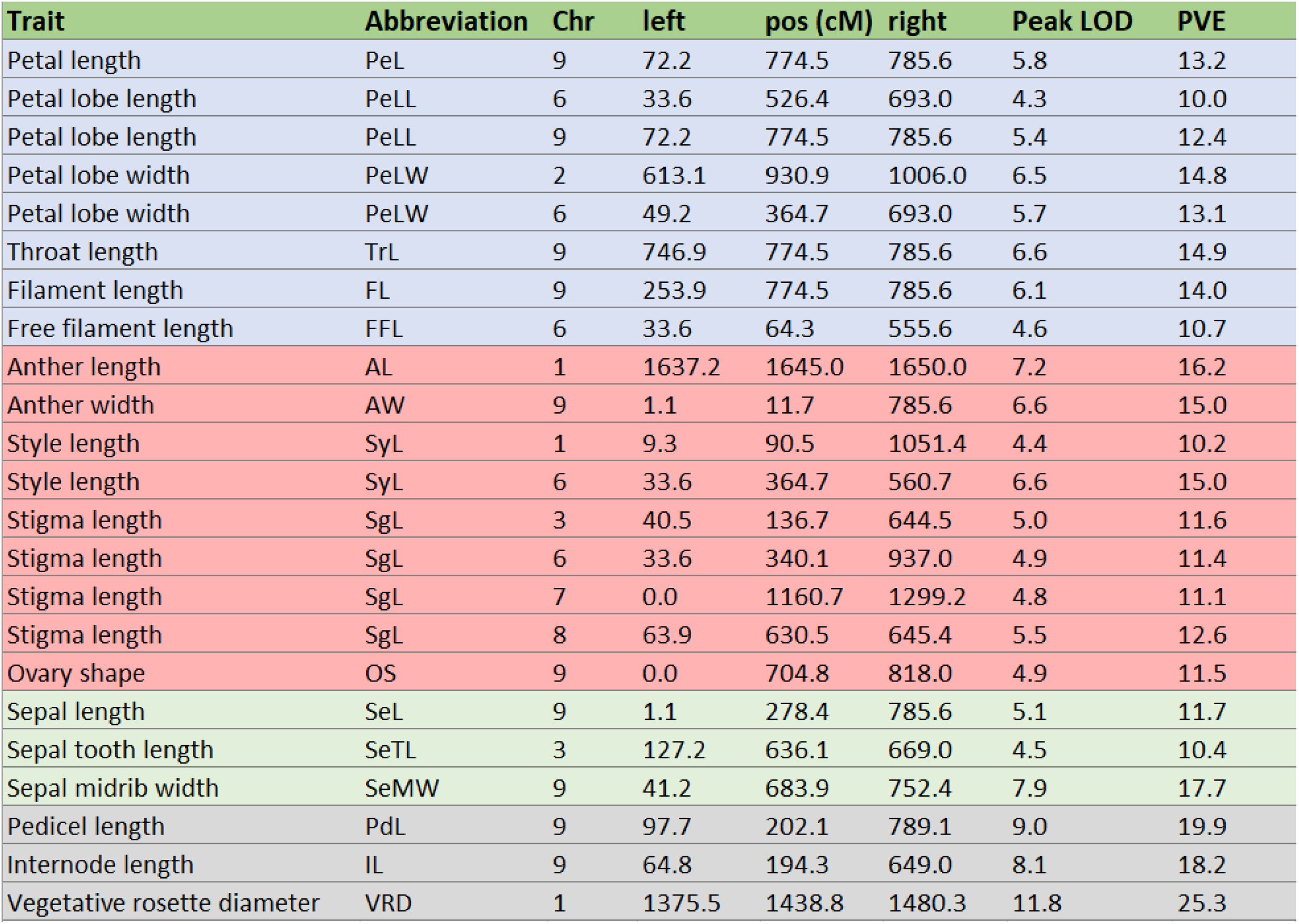
List of traits with significant QTL. Intervals were calculated using a 1.5 LOD drop. Left and right endpoints, position of the highest-correlated marker, and Percent Variance Explained (PVE) are shown.

| Trait | Abbreviation | Chr | left | pos (cM) | right | Peak LOD | PVE |
| --- | --- | --- | --- | --- | --- | --- | --- |
| Petal length | PeL | 9 | 72.2 | 774.5 | 785.6 | 5.8 | 13.2 |
| Petal lobe length | PeLL | 6 | 33.6 | 526.4 | 693.0 | 4.3 | 10.0 |
| Petal lobe length | PeLL | 9 | 72.2 | 774.5 | 785.6 | 5.4 | 12.4 |
| Petal lobe width | PeLW | 2 | 613.1 | 930.9 | 1006.0 | 6.5 | 14.8 |
| Petal lobe width | PeLW | 6 | 49.2 | 364.7 | 693.0 | 5.7 | 13.1 |
| Throat length | TrL | 9 | 746.9 | 774.5 | 785.6 | 6.6 | 14.9 |
| Filament length | FL | 9 | 253.9 | 774.5 | 785.6 | 6.1 | 14.0 |
| Free filament length | FFL | 6 | 33.6 | 64.3 | 555.6 | 4.6 | 10.7 |
| Anther length | AL | 1 | 1637.2 | 1645.0 | 1650.0 | 7.2 | 16.2 |
| Anther width | AW | 9 | 1.1 | 11.7 | 785.6 | 6.6 | 15.0 |
| Style length | SyL | 1 | 9.3 | 90.5 | 1051.4 | 4.4 | 10.2 |
| Style length | SyL | 6 | 33.6 | 364.7 | 560.7 | 6.6 | 15.0 |
| Stigma length | SgL | 3 | 40.5 | 136.7 | 644.5 | 5.0 | 11.6 |
| Stigma length | SgL | 6 | 33.6 | 340.1 | 937.0 | 4.9 | 11.4 |
| Stigma length | SgL | 7 | 0.0 | 1160.7 | 1299.2 | 4.8 | 11.1 |
| Stigma length | SgL | 8 | 63.9 | 630.5 | 645.4 | 5.5 | 12.6 |
| Ovary shape | OS | 9 | 0.0 | 704.8 | 818.0 | 4.9 | 11.5 |
| Sepal length | SeL | 9 | 1.1 | 278.4 | 785.6 | 5.1 | 11.7 |
| Sepal tooth length | SeTL | 3 | 127.2 | 636.1 | 669.0 | 4.5 | 10.4 |
| Sepal midrib width | SeMW | 9 | 41.2 | 683.9 | 752.4 | 7.9 | 17.7 |
| Pedicel length | PdL | 9 | 97.7 | 202.1 | 789.1 | 9.0 | 19.9 |
| Internode length | IL | 9 | 64.8 | 194.3 | 649.0 | 8.1 | 18.2 |
| Vegetative rosette diameter | VRD | 1 | 1375.5 | 1438.8 | 1480.3 | 11.8 | 25.3 |

**Figure 6.**
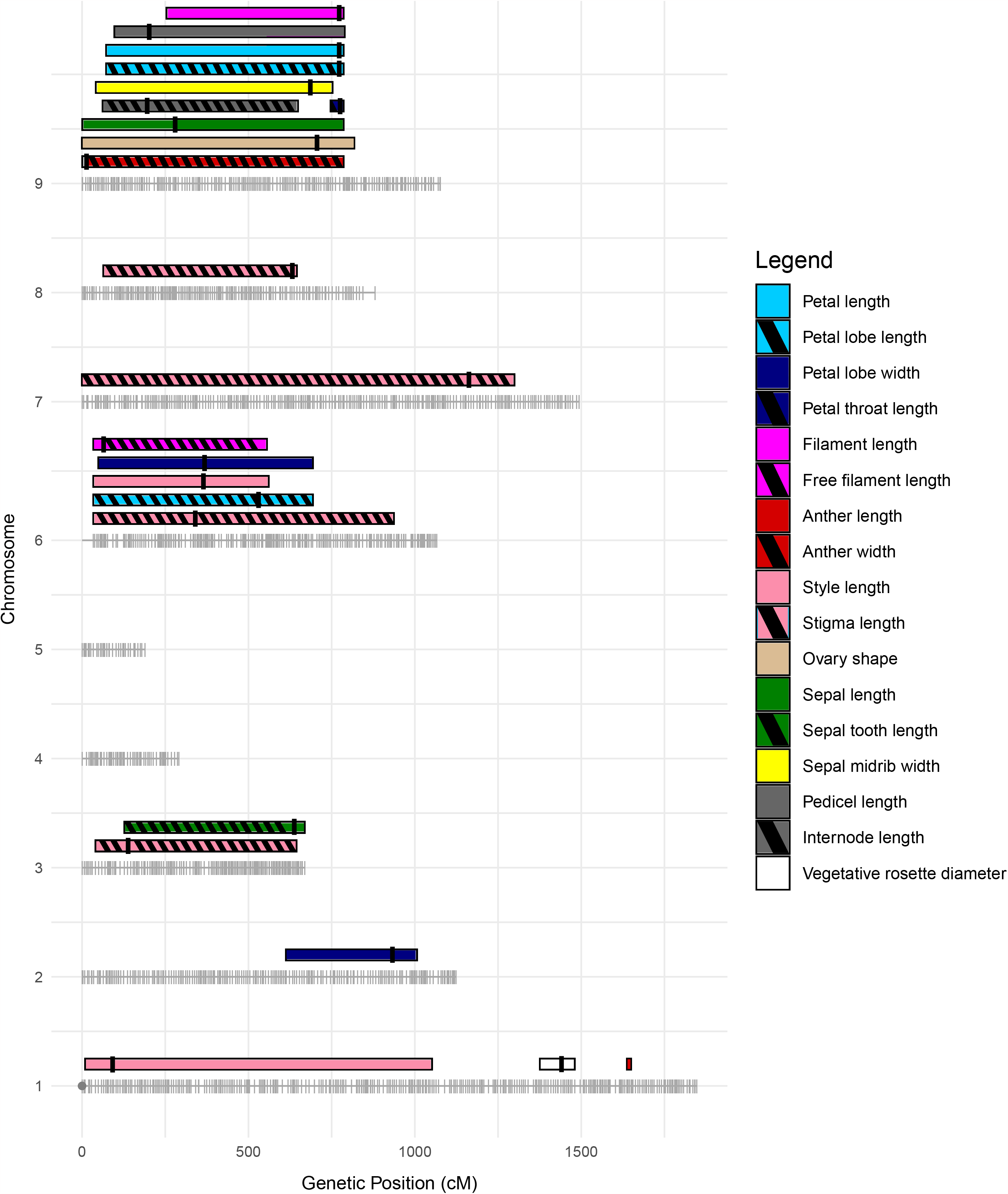
QTL intervals for all significant traits. Marker distribution is shown in gray along each chromosome. Rectangles represent 1.5 LOD drop intervals with right and left endpoints shown. The line within each rectangle represents the position of the highest-correlated marker.

Notably, 10 of the 23 QTL localized to Chromosome 9, including the PeL, PeLL, TrL, FL, AW, OS, SeL, SeMW, PdL, and IL traits (Table 3). Nine of these QTL have a width greater than 500 cM, making it difficult to determine if the colocalization of these traits is due to a single pleiotropic locus or multiple linked loci.

In addition to the QTL hotspot on Chromosome 9, four QTL mapped to similar regions on Chromosome 6. These QTL correspond to PeLL, PeLW, SyL, and SgL. Only one of these traits (PeLL) overlaps with the multiple QTL found on Chromosome 9. As before, this may indicate a single pleiotropic locus or multiple linked loci.

Most of the QTL identified had an effect of 10-15% of variance (PvE) explained in the F2 population. The three exceptions to this are vegetative rosette diameter (25.3%), sepal midrib width (17.7%), and anther length (16.2%) (Table 3). VRD shows evidence of a relatively large-effect QTL localized to a genetically narrow region (<200cM) on Chromosome 1. AL also localizes to a narrow QTL, with a smaller effect on phenotype.

## Discussion

In this study, we have described floral trait averages, correlations, and QTLs from an F2 population derived from a cross between *G. yorkii* and *G. capitata*. These two species are divergent for floral color and inflorescence architecture, which have been described in a previous publication (Jarvis *et al*. 2022). Here, we explore the genetic structure of quantitative floral traits that distinguish *G. yorkii* and *G. capitata*. Several trait QTLs occupy distinct regions of the genome, which are good candidates for QTLs that control only a single trait. For example, SgL on chromosomes 7 and 8, PeLW on chromosome 2, and VRD on chromosome 1 all have QTLs that don’t colocalize with other traits. On the other hand, multiple QTLs colocalized in hotspots on Chromosomes 6 and 9. Similar QTL studies in other plant species often find regions with colocalizing QTLs (Bouck *et al*. 2007; Zhu *et al*. 2014; Wessinger *et al*. 2014; Kostyun *et al*. 2019). This shared genetic basis can be due either to a) multiple linked loci that are difficult to separate via recombination but control traits independently or b) pleiotropic loci that simultaneously control several traits (Lande 1984). Though this study does not have the resolution to distinguish multiple linked loci from pleiotropic loci, future work with advanced generation mapping lines including random intermated and/or recombinant inbred lines may help resolve multilocus from pleiotropic QTL (Conner 2002; Mackay 2003). Regardless of the underlying cause, it is clear that multiple trait QTLs overlap on chromosomes 3, 6, and 9. In particular, chromosome 9 has a high concentration of overlapping trait QTLs, including PeL, PeLL, TrL, FL, AW, OS, SeL, SeMW, PdL, and IL. Because of the broad QTL intervals, it is not yet possible to begin considering candidate genes for these traits. However, this region explains up to 20% of the variance for 10 floral traits, making it the most influential QTL region identified in this study.

The co-localization of QTLs is consistent with the morphological correlations we observe between traits. Within the F2 population, there were two clear groups of strongly correlated traits: one consisting of corolla and style traits and the other consisting of sepal traits. Interestingly, some traits have a clear lack of correlation with any other traits. AL, despite having a strong correlation with AW, shows only weak correlations with other corolla traits. This contrasts with reported studies in *Mimulus* (Fenster and Ritland 1994; Fishman *et al*. 2002; Hall *et al*. 2006; Fishman *et al*. 2015) that have higher correlations of AL with other corolla traits, suggesting a weaker connection of AL and corolla traits within *Gilia* species. VRD shows strikingly low correlations with all other traits, showing that floral trait correlations are not biased by the plant’s overall size. SgL is previously known to be connected to calyx pubescence, calyx reflexion, and capsule dehiscence in intraspecies crosses between *G. capitata* subspecies (Grant 1950). Our results agree with that study, in that SgL shows very weak associations with all other floral traits such as PeLW. In summary, most of the corolla and sepal traits correlate strongly within, but not across, their respective floral whorls. Some traits, like AL, do not seem to be connected directly to other traits, even with physically adjacent traits.

While many traits showed high heritabilities between the parent and F1 generations (Table 1), the variance explained by the discovered QTLs (Table 3) is significantly lower. For example, FFL shows a heritability of 0.93, while the single QTL discovered for this trait explains only 10.7% of the variance within the F2 population. This discrepancy may be due to physical chromosomal regions not included in the genetic map, especially on chromosomes 4 and 5.

Any causative QTLs located within these regions are not detectable by our QTL analysis. Another possibility is that some of the trait variation consists of small-effect loci that don’t pass the significance threshold. Some traits have significantly more agreement between their heritability and percent variance explained. For example, VRD has a heritability of 0.18 and its single QTL explains 25.3% of the F2 variation, indicating that the genetic component of this trait is fully captured by that QTL. Similarly, SgL has a heritability of 0.42 and has four QTL that, when combined, explain 46.7% of the F2 variation. In summary, most of the traits examined appear to have genetic variation beyond that explained by the discovered QTLs, whereas two traits, VRD and SgL, appear to be explained entirely by their respective QTLs.

The adaptive purpose, if indeed one exists, for the morphological differences of *G. yorkii* and *G. capitata* flowers is still unclear. Considering the importance of the size and shape of floral organs for successful pollination, it seems likely that there may be pollinator differences between these species. Verne Grant documented the potential pollinators of *G. capitata* and some of its subspecies (Grant and Grant 1965), which attract a wide range of insect visitors, including various bees, beeflies, beetles, and butterflies (Grant and Grant 1965). Since the discovery of *G. yorkii*, its pollinators have not yet been reported. Comparing *G. yorkii* to other species within Polemoniaceae that share a similar floral shape and inflorescence structure, such as *G. achilleifolia ssp*.*multicaulis, G. angelensis*, and *G. tricolor*, it appears likely that pollinator classes are similar to *G. capitata* (i.e. bees, beeflies, beetles, and butterflies), but may be smaller in size overall (Grant and Grant 1965). Overall, the reduction in stamen exsertion and lack of pigmentation in *G. yorkii* could be consistent with some level of self-pollination, although the relative increase in petal lobe size may serve to compensate as an attractant for insect pollinators. Field observations are needed to verify these suspected pollinator attraction and reproductive system differences.

Overall, we have presented the first study connecting the floral traits of two *Gilia* species to their genetic underpinnings. These traits are confirmed to be quantitative in nature, as evidenced by the absence of major effect QTL explaining more than 50% of phenotypic variation, and many traits appear to co-localize to similar regions in the genome. Along with this, petal and sepal traits constitute two distinct groups of correlated traits, suggesting that each group shares pleiotropic or linked genes that control sets of traits together. Future work may tease apart these possibilities through fine-mapping approaches with greater recombination frequencies than is possible in an F2 intercross.

## Data Availability

All phenotypic and genotypic data used for the QTL analysis are available in supplementary material. In addition, all data analysis scripts are available on Github at https://github.com/detemplej/Gilia-QTL-data. Data for the morphometric analysis and the original microscope images of flowers are available upon request.

## Funding

Support for this research was generously provided by the BYU Department of Biology to CW as well as a College Undergraduate Research Award (CURA) from the BYU College of Life Sciences to JD.

## Conflicts of Interest

The author(s) declare no conflict of interest.

## Literature cited

Ballerini ES, Min Y, Edwards MB, Kramer EM, Hodges SA. 2020. Popovich, encoding a c2h2 zinc-finger transcription factor, plays a central role in the development of a key innovation, floral nectar spurs, in aquilegia. Proceedings of the National Academy of Sciences of the United States of America. 117:22552–22560.

Bouck A, Wessler SR, Arnold ML. 2007. Qtl analysis of floral traits in louisiana iris hybrids. Evolution. 61:2308–2319.

Bradshaw HD, Wilbert SM, Otto KG, Schemske DW. 1995. Genetic mapping of floral traits associated with reproductive isolation in monkeyflowers (mimulus). Nature 1995 376:6543. 376:762–765.

Broman KW, Sen S. 2009. Statistics for biology and health a guide to qtl mapping with r/qtl. .

Broman KW, Wu H, Sáunak Sen, Churchill GA. 2003. R/qtl: Qtl mapping in experimental crosses. Bioinformatics. 19:889–890.

Brothers AN, Barb JG, Ballerini ES, Drury DW, Knapp SJ, Arnold ML. 2013. Genetic architecture of floral traits in iris hexagona and iris fulva. Journal of Heredity. 104:853–861.

Campitelli BE, Kenney AM, Hopkins R, Soule J, Lovell JT, Juenger TE. 2018. Genetic mapping reveals an anthocyanin biosynthesis pathway gene potentially influencing evolutionary divergence between two subspecies of scarlet gilia (ipomopsis aggregata). Molecular Biology and Evolution. 35:807–822.

Chen YY, Nishii K, Kidner C, Hackett CA, Möller M. 2020. Qtl dissection of floral traits in streptocarpus (gesneriaceae). Euphytica. 216:1–20.

Conner JK. 2002. Genetic mechanisms of floral trait correlations in a natural population. Nature 2002 420:6914. 420:407–410.

Eckert CG, Manicacci D, Barrett SCH. 1996. Genetic drift and founder effect in native versus introduced populations of an invading plant, lythrum salicaria (lythraceae). Evolution. 50:1512.

Fenster CB, Ritland K. 1994. Quantitative genetics of mating system divergence in the yellow monkeyflower species complex. Heredity 1994 73:4. 73:422–435.

Fishman L, Beardsley PM, Stathos A, Williams CF, Hill JP. 2015. The genetic architecture of traits associated with the evolution of self-pollination in mimulus. New Phytologist. 205:907–917.

Fishman L, Kelly AJ, Willis JH. 2002. Minor quantitative trait loci underlie floral traits associated with mating system divergence in mimulus. Evolution. 56:2138–2155.

Freeling M. 2001. Grasses as a single genetic system. reassessment 2001. Plant Physiology. 125:1191–1197.

Goh L, Yap VB. 2009. Effects of normalization on quantitative traits in association test. BMC Bioinformatics. 10:1–8.

Goodwillie C, Ritland C, Ritland K. 2006. The genetic basis of floral traits associated with mating system evolution in leptosiphon (polemoniaceae): An analysis of quantitative trait loci. Evolution. 60:491–504.

Grant V. 1949. Seed germination in gilia capitata and its relatives on jstor.

Grant V. 1950. Genetic and taxonomic studies in gilia: I. gilia capitata. Aliso: A Journal of Systematic and Floristic Botany. 2:239–316.

Grant V. 1966. The selective origin of incompatibility barriers in the plant genus gilia. 10.1086/282404. 100:99–118.

Grant V, Grant KA. 1965. Flower Pollination in the Phlox Family. Columbia University Press.

Hall MC, Basten CJ, Willis JH. 2006. Pleiotropic quantitative trait loci contribute to population divergence in traits associated with 10 Interspecies floral morphology QTL in Gilia life-history variation in mimulus guttatus. Genetics. 172:1829–1844.

Hodges SA, Whittall JB, Fulton M, Yang JY. 2002. Genetics of floral traits influencing reproductive isolation between aquilegia formosa and aquilegia pubescens. The American Naturalist. 159:S51–S60.

Jarvis DE, Maughan PJ, DeTemple J, Mosquera V, Li Z, Barker MS, Johnson LA, Whipple CJ. 2022. Chromosome-scale genome assembly of gilia yorkii enables genetic mapping of floral traits in an interspecies cross. Genome Biology and Evolution. 14.

Johnson LA, Porter JM. 2017. Fates of angiosperm species following long-distance dispersal: Examples from american amphitropical polemoniaceae. American Journal of Botany. 104:1729–1744.

Kostyun JL, Gibson MJ, King CM, Moyle LC. 2019. A simple genetic architecture and low constraint allow rapid floral evolution in a diverse and recently radiating plant genus. New Phytologist. 223:1009–1022.

Lande R. 1984. The genetic correlation between characters maintained by selection, linkage and inbreeding. Genetics Research. 44:309–320.

Liang M, Chen W, LaFountain AM, Liu Y, Peng F, Xia R, Bradshaw HD, Yuan YW. 2023. Taxon-specific, phased sirnas underlie a speciation locus in monkeyflowers. Science. 379:576–582.

Mackay TF. 2003. The genetic architecture of quantitative traits. 10.1146/annurev.genet.35.102401.090633. 35:303–339.

Mertens A, Brys R, Schouppe D, Jacquemyn H. 2018. The impact of floral morphology on genetic differentiation in two closely related biennial plant species. AoB Plants. 10.

Miles CM, Wayne M. 2008. Quantitative trait locus (qtl) analysis | learn science at scitable.

Pinheiro F, Barros FD, Palma-Silva C, Meyer D, Fay MF, Suzuki RM, Lexer C, Cozzolino S. 2010. Hybridization and introgression across different ploidy levels in the neotropical orchids epidendrum fulgens and e. puniceoluteum (orchidaceae). Molecular Ecology. 19:3981–3994.

Porter JM. 2012. Gilia, in jepson flora project.

Rieseberg LH, Wood TE, Baack EJ. 2006. The nature of plant species. Nature 2005 440:7083. 440:524–527.

Roux C, Pannell JR. 2019. The opposing effects of genetic drift and haldane’s sieve on floral-morph frequencies in tristylous metapopulations. The New Phytologist. 224:1229.

Sapir Y, Gallagher MK, Senden E. 2021. What maintains flower colour variation within populations? Trends in Ecology & Evolution. 36:507–519.

Shevock JR, Day AG. 1998. A new gilia (polemoniaceae) from limestone outcrops in the southern sierra nevada of california. Madroño. 45:137–140.

Tremblay RL, Ackerman JD. 2001. Gene flow and effective population size in lepanthes (orchidaceae): a case for genetic drift. Biological Journal of the Linnean Society. 72:47–62.

Wessinger CA, Hileman LC, Rausher MD. 2014. Identification of major quantitative trait loci underlying floral pollination syndrome divergence in penstemon. Philosophical Transactions of the Royal Society B: Biological Sciences. 369.

Yoshida Y, Honjo M, Kitamoto N, Ohsawa R. 2008. Genetic variation and differentiation of floral morphology in wild primula sieboldii evaluated by image analysis data and ssr markers. Breeding Science. 58:301–307.

Zhu R, Gao Y, Zhang Q. 2014. Quantitative trait locus mapping of floral and related traits using an f2 population of aquilegia. Plant Breeding. 133:153–161.

